# Genetic dissection of Down syndrome-associated alterations in APP/amyloid-β biology using mouse models

**DOI:** 10.1101/2020.06.19.162115

**Authors:** Justin L. Tosh, Ellie Rhymes, Paige Mumford, Heather T. Whittaker, Laura J. Pulford, Sue J. Noy, Karen Cleverley, Matthew C. Walker, Victor L.J. Tybulewicz, Rob C. Wykes, Elizabeth M.C Fisher, Frances K. Wiseman

## Abstract

Individuals who have Down syndrome (caused by trisomy of chromosome 21), have a greatly elevated risk of early-onset Alzheimer’s disease, in which amyloid-β accumulates in the brain. Amyloid-β is a product of the chromosome 21 gene *APP* (amyloid precursor protein) and the extra copy or ‘dose’ of *APP* is thought to be the cause of this early-onset Alzheimer’s disease. However, other chromosome 21 genes likely modulate disease when in three-copies in people with Down syndrome. Here we show that an extra copy of chromosome 21 genes, other than *APP*, influences APP/Aβ biology. We crossed Down syndrome mouse models with partial trisomies, to an *APP* transgenic model and found that extra copies of subgroups of chromosome 21 gene(s) modulate amyloid-β aggregation and *APP* transgene-associated mortality, independently of changing amyloid precursor protein abundance. Thus, genes on chromosome 21, other than *APP*, likely modulate Alzheimer’s disease in people who have Down syndrome.

## INTRODUCTION

Down syndrome (DS), which occurs in approximately 1 in 1000 births, is the most common cause of early-onset Alzheimer’s disease-dementia (AD-DS) (*1*). Approximately 6 million people have DS world-wide and by the age of 65 two-thirds of these individuals will have a clinical dementia diagnosis. Moreover, approaching 100% of women with DS will have a dementia diagnosis by the age of 80 (*2*) and dementia is now a leading cause of death for people who have DS in the UK (*3*). Trisomy of human chromosome 21 (Hsa21) and the resulting abnormal gene dosage also significantly affect neurodevelopment, neuronal function and other aspects of physiology, such as cardiovascular and immune systems, giving rise to the spectrum of features seen in DS (*4*).

Clinical-genetic studies demonstrate that the Hsa21 gene amyloid precursor protein (*APP)* is central to early AD onset in people with DS (*5, 6*), but other genes, including some found on Hsa21, modulate age of dementia onset (*7*–*9*). APP undergoes enzymatic cleavage within the brain to form numerous fragments, including amyloid-β which accumulates in AD-associated amyloid plaques, and C-terminal fragments (CTF) that may impair intracellular processes (*10*). However, how trisomy of Hsa21 genes, other than *APP*, impacts APP biology and subsequent neurodegeneration and dementia is not well understood (*1*).

Individual DS phenotypes likely arise from an extra copy of specific genes on Hsa21, which is currently estimated to carry 234 protein-coding genes (*11*). While not all Hsa21 genes are ‘dosage-sensitive’, the increased dosage of some genes may lead to increased transcript and protein levels that result in biologically relevant molecular and cellular changes. Identification of the Hsa21 genes that affect AD-dementia development will further understanding of AD-DS mechanisms and may also provide novel insight into neurodegeneration in the euploid population by the identification of key pathways. Moreover, understanding the development of AD-DS is of critical importance for the translation of AD prevention therapies for people with DS.

In a previous study we crossed a mouse model of DS (the Tc1 mouse) that does not have an additional copy of *APP* to a transgenic mouse overexpressing APP with AD-causing mutations (‘J20’ mice). This work demonstrated that an additional copy of Hsa21, independent of an extra copy of *APP*, exacerbated amyloid-β aggregation and deposition, enhanced *APP* transgene-associated mortality, altered behaviour and reduced cognitive performance in double mutant progeny that model AD-DS (*12*). In the same system we also observed that an additional copy of Hsa21, independently of an extra copy of *APP*, increased APP-CTF fragments in male but not female brains in AD-DS mice.

Here using independent DS mouse models (*13, 14*) we investigated these phenotypes further and determined that a causal gene(s) for elevated amyloid-β aggregation lies between *Mir802* and *Zbtb21* (within the ‘Dp(16)3Tyb’ mouse duplication). We also found that a dosage sensitive gene(s) that enhances *APP* transgene-associated mortality is located between *Mis18a* and *Runx1* (within the ‘Dp(16)2Tyb’ mouse duplication) and that dosage-sensitive genes that protect against *APP* transgene-associated mortality are located between *Mir802* and *Zbtb21* (Dp(16)3Tyb duplication), *Prmt2* and *Pdxk* (Dp(17)1Yey duplication) and also between *Abcg1* and *Rrp1b* (Dp(10)1Yey duplication). We went on to show that the rescue of mortality by the Dp(10)1Yey duplication occurred independently of changes to the frequency or duration of *APP* transgene-associated seizures.

These data show that an extra copy of multiple chromosome 21 gene orthologues modulate multiple AD-related phenotypes in mouse models. Similar mechanisms may also occur in people who have DS when they develop AD-dementia and may contribute to the AD-clinical differences that occur within this population.

## RESULTS

### Survival of tgAPP mouse is modulated by additional copies of chromosome 21 mouse homologues

To determine whether an additional copy of chromosome 21 genes other than *APP* modified APP/amyloid-β biology we crossed the J20 *APP* transgenic (tgAPP) mouse model, which expresses human APP with AD-associated point mutations, with a panel of mouse models of DS (**Fig.1A**). These DS mouse models have segmental duplications of defined regions of the mouse genome that are syntenic with Hsa21. Hsa21 has homology to three regions of the mouse genome, within mouse chromosomes 10, 16 and 17 (Mmu10, Mmu16, Mmu17). Each model has an extra copy of a subset of Hsa21-orthologous genes and can be used to determine which Hsa21 genes cause DS-associated phenotypes (*13, 15*). We systematically assessed crosses of these DS models with the J20 tgAPP mouse to determine how each segmental duplication affected tgAPP-associated phenotypes. Mice with the tgAPP transgene and a segmental duplication were compared to littermates that only carried tgAPP to assess the effect of the segmental duplication on APP/Aβ biology (**Fig. 1B**).

**Fig. 1.**
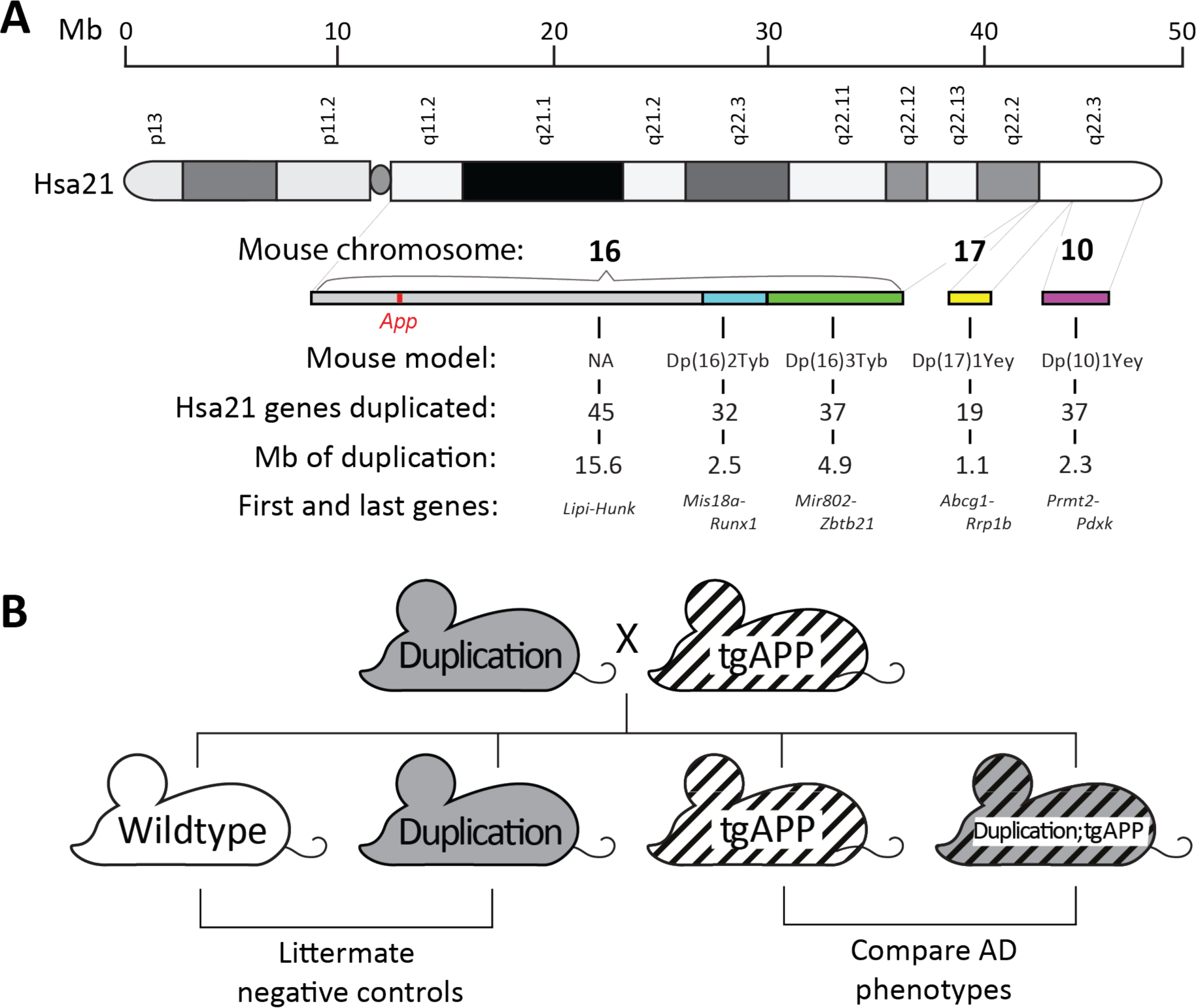
Mapping DS-AD associated phenotypes: analysis of progeny from crosses of J20 tgAPP with DS segmental duplication models. **(A)** Regions of Hsa21 homology on mouse chromosomes 16, 17 and 10 in the Dp(16)2Tyb, Dp(16)3Tyb, Dp(10)1Yey and Dp(17)1Yey segmental duplication models of DS. An ideogram of Hsa21 with major karyotypic bands is shown, with a megabase (Mb) scale. Below this are graphical representations of the relative sizes of Hsa21 orthologous regions in the mouse and the subregions which are segmentally duplicated in each DS mouse model used in this study. These are colour coded: Dp(16)2Tyb is light blue, Dp(16)3Tyb is green, Dp(10)1Yey is purple, and Dp(17)1Yey is yellow. “NA” refers to a subregion not modelled in this study. The number of Hsa21 orthologous genes, Mb size, first and last genes in each duplication are shown. **(B)** Schematic of the cross of the segmental duplication models of DS crossed with the J20 *APP* transgenic model (tgAPP).

*APP* transgenic mice often exhibit elevated mortality, likely because of the epileptogenic effect of APP or an APP cleavage product (*16*). We determined whether mortality in our J20 tgAPP progeny was modulated by the extra copy of Hsa21 mouse homologues in the DS mouse mapping panel (**Fig. 2**), because we had determined previously that trisomy of human chromosome 21 elevated tgAPP-associated mortality in another mouse cross (*12*). We found an additional copy of a gene(s) between *Mis18a* and *Runx1* (Dp(16)2Tyb segmental duplication on Mmu16) significantly reduced survival associated with J20 tgAPP prior to 6-months of age (**Fig. 2A**). Thus, for animal welfare we ceased this experimental cross and only a 6-month of age time-point was produced, and we did not investigate phenotypes in older animals.

**Fig. 2.**
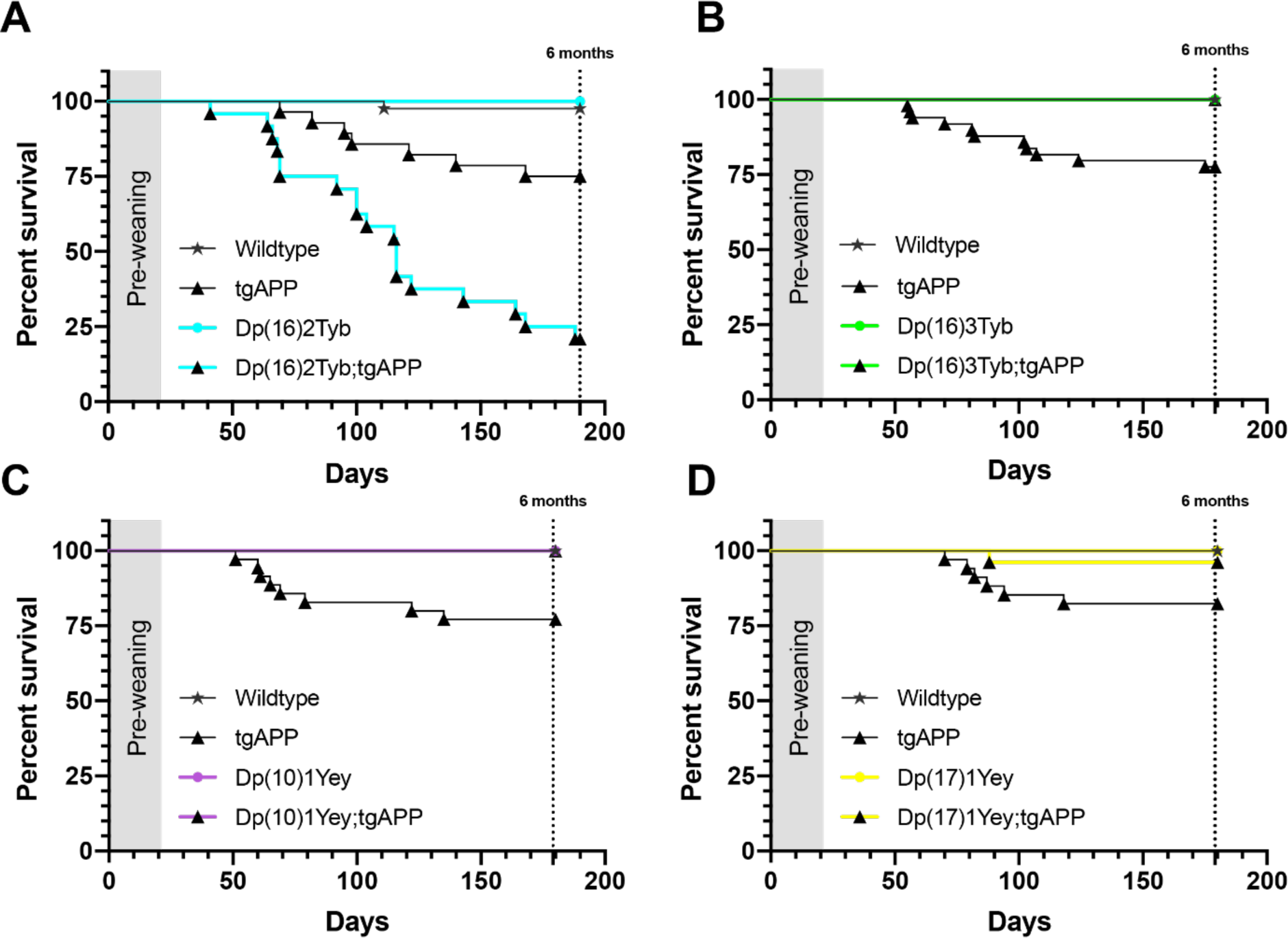
The effect of DS segmental duplication models on J20 tgAPP-associated mortality. **(A)** Survival of Dp(16)2Tyb;tgAPP male and female mice to 6-months of age is significantly reduced compared to tgAPP controls (Mantel-Cox log-rank test X^2^ = 47.872 p <0.001). (Wildtype male n = 21, female n = 16; Dp(16)2Tyb male n = 7, female n = 6; tgAPP male n = 16, female n = 12; Dp(16)2Tyb;tgAPP male n = 7, female n = 21). **(B)** Survival of Dp(16)3Tyb;tgAPP male and female mice to 6-months of age is significantly increased compared to tgAPP controls (Mantel-Cox log-rank test X^2^ = 33.58 p <0.001). (Wildtype male n = 23, female n = 34; Dp(16)3Tyb male n = 19, female n = 20; tgAPP male n = 21, female n = 28; Dp(16)3Tyb;tgAPP male n = 18, female n = 20). **(C)** Survival of Dp(10)1Yey;tgAPP male and female mice to 6-months of age is significantly increased compared to tgAPP controls (Mantel-Cox log-rank test X^2^ = 8.414 p = 0.004). (Wildtype male n = 15, female n = 23; Dp(10)1Yey male n = 19, female n = 27; tgAPP male n = 17, female n = 18; Dp(10)1Yey;tgAPP male n = 11, female n = 22). **(D)** Survival of Dp(17)1Yey;tgAPP male and female mice to 6-months of age is significantly increased compared to tgAPP controls (Mantel-Cox log-rank test X^2^ = 12.56 p = 0.0057). (Wildtype male n = 14, female n = 18; Dp(10)1Yey male n = 11, female n = 15; tgAPP male n = 16, female n = 18; Dp(10)1Yey;tgAPP male n = 12, female n = 14). Both sexes included in analysis.

Conversely an additional copy of the regions between *Mir802* and *Zbtb21* (Dp(16)3Tyb duplication on Mmu16) (**Fig. 2B**), *Prmt2* and *Pdxk* (Dp(10)1Yey duplication on Mmu10) (**Fig. 2C**) and *Abcg1* and *Rrp1b* (Dp(17)1Yey duplication on Mmu17) (**Fig. 2D**) all partially rescued J20 tgAPP-associated reduced survival.

These data suggest that an additional copy of at least four genes on Hsa21 modulates tgAPP-related mortality. Moreover, that an extra copy of genes on Hsa21 both exacerbate and alleviate tgAPP-associated mortality. This may be the result of suppression of tgAPP-associated seizures which are thought to be the major cause of elevated mortality in the J20 mouse model. Alternatively, seizures may still occur but the mice may be protected from post-seizure mortality. Seizure occurrence is linked to the abundance of APP and its cleavage fragments (*16, 17*), thus changes to mortality may result from alterations in these proteins. Thus, we next tested these potential mechanisms.

### Segmental duplications do not alter FL-APP abundance but the Dp(16)3Tyb duplication causes an increase in cortical α-CTF

The observed alteration in tgAPP-associated mortality may result from changes to the abundance of full-length APP (FL-APP) or one of its cleavage fragments (*16, 18, 19*). Thus, we investigated the abundance of full-length APP (FL-APP) in the progeny of a cross of Dp(16)3Tyb, Dp(10)1Yey or Dp(17)1Yey duplications with tgAPP J20 mice at 3-months of age. We found no evidence that abundance of FL-APP (**Fig. S1A**) was altered by an additional copy of the Dp(16)3Tyb, Dp(17)1Yey or Dp(10)1Yey regions, in the cortex. We were not able to investigate the effect of the Dp(16)2Tyb region *in vivo* on FL-APP abundance because the elevated mortality observed in the intercross prevented the ethical generation of tissue samples for these experiments.

Using an alternative mouse model of DS (the ‘Tc1’ mouse) we have previously shown that an extra copy of Hsa21 raises the abundance of alpha and beta C-terminal fragments of APP (α-CTF and β-CTF) in the brain of male mice (*12*). Here we show that an additional copy of a gene(s) between *Mir802* and *Zbtb21* (Dp(16)3Tyb segmental duplication) is sufficient to raise the level of α-CTF in the cortex in male and female mice (**Fig. 3A**). No significant changes in β-CTF levels were detected in the presence of the Dp(16)3Tyb duplication (**Fig. 3A**). α-CTF and β-CTF levels were not altered in the Dp(10)1Yey or Dp(17)1Yey duplication models (**Fig. 3B, C**). We were not able to investigate the effect of the Dp(16)2Tyb region *in vivo* on α-CTF and β-CTF abundance because the elevated mortality observed in the intercross prevented the ethical generation of tissue samples for these experiments.

**Fig. 3.**
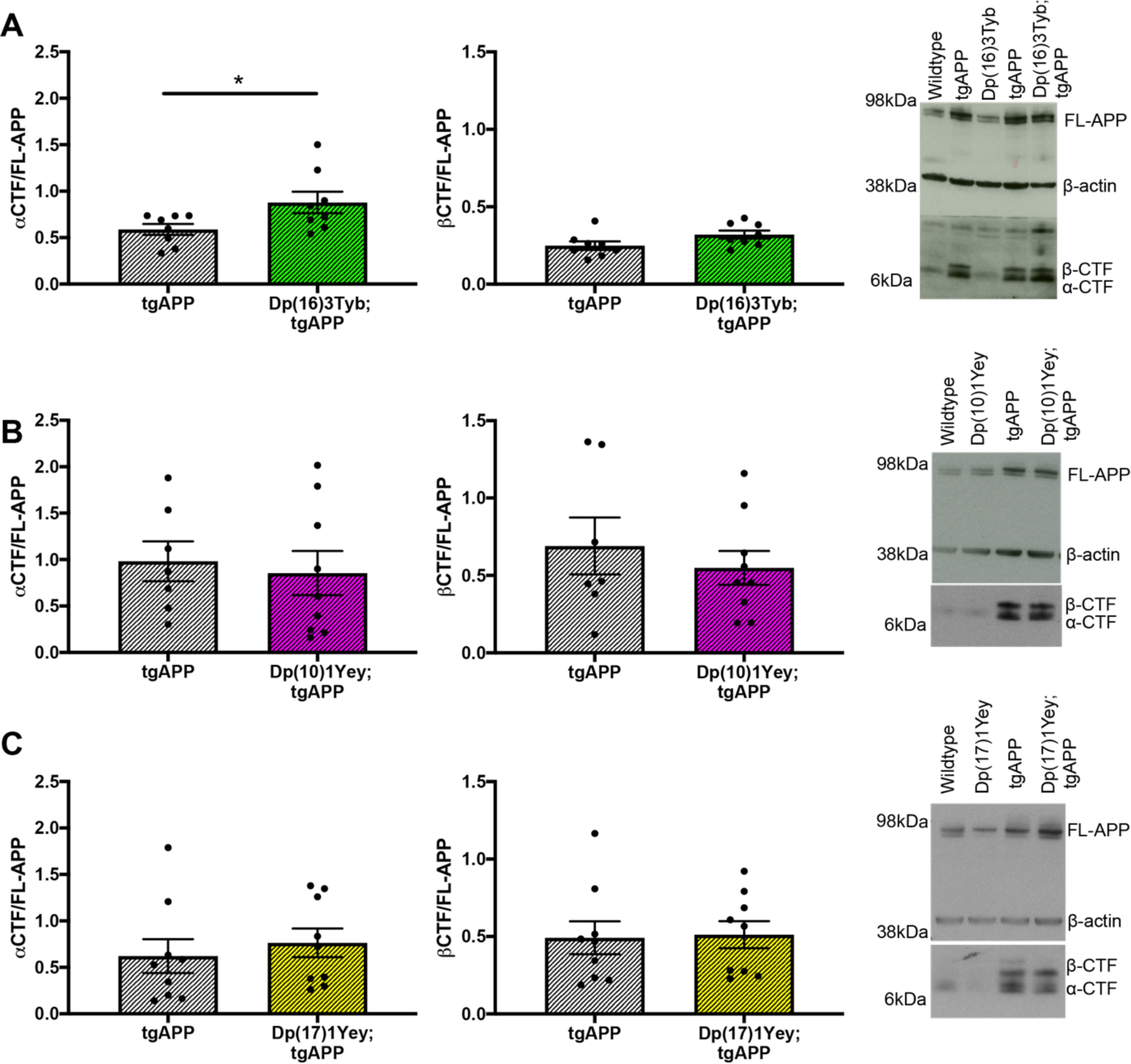
The effect of DS segmental duplication models on CTF abundance in the brain. **(A-C)** The relative abundance of APP β-C-terminal fragment (β-CTF) and APP α-C-terminal fragment (α-CTF) compared to full-length APP (FL-APP) was measured by western blot using A8717 primary antibody in the cortex at 3-months of age in female and male mice. **(A)** Significantly more α-CTF was observed in Dp(16)3Tyb;tgAPP (n = 8, 4 male and 4 female) than tgAPP (n = 8, 4 male and 4 female) controls (F(1,12) = 5.226, p = 0.041), β-CTF was not raised in this double mutant mouse (F(1,13) = 3.005, p = 0.107). **(B)** In Dp(10)1Yey;tgAPP mice (n = 11, 7 male and 4 female) neither α-CTF (F(1,12) = 0.143, p = 0.712) nor β-CTF (F(1,12) = 0.291, p = 0.599) abundance differed from tgAPP (n = 7, 3 male and 4 female) controls. **(C)** In Dp(17)1Yey;tgAPP (n = 9, 6 male and 3 female) mice neither α-CTF (F(1,13) = 0.350, p = 0.363) nor β-CTF (F(1,13) = 0.566, p = 0.465) abundance differed from tgAPP (n = 9, 3 male and 6 female) controls. Error bars show SEM, data points are independent mice.

To further investigate the effect of the Dp(16)3Tyb duplication on α-CTF and β-CTF abundance we measured levels in the hippocampus at 3-months of age; no change in FL-APP, α-CTF or β-CTF was observed (**Fig. S2**). Therefore, the rescue of tgAPP-associated mortality in the Dp(16)3Tyb and Dp(10)1Yey mouse models of DS occurs independently of a change in FL-APP or β-CTF in two key brain regions in the J20 tgAPP model. Moreover, the increase in α-CTF abundance in the Dp(16)3Tyb segmental duplication model is brain region-dependent but sex-independent in contrast to our previous work in the Tc1 DS model.

### Additional copies of Hsa21 homologues modulate aggregation of amyloid-β in the brain

Raised amyloid-β may cause excitotoxicity and seizure related mortality (*18*). Moreover, our previous study using the Tc1 mouse model of trisomy of chromosome 21 demonstrated that aggregation of amyloid-β_42_ is increased by the additional chromosome, independently of an additional copy of *APP*. Thus, we determined if an additional copy of the mouse Hsa21 orthologues altered amyloid-β biology. We fractionated soluble and insoluble (aggregated) cortical proteins and quantified levels of amyloid-β_40_ and amyloid-β_42_.

The abundance of insoluble amyloid-β_42_ in the cortex was decreased in Dp(16)2Tyb;tgAPP mice compared to tgAPP littermates at 6-months of age in both males and females (**Fig. 4A**). However, given the high mortality of tgAPP;Dp(16)2Tyb mice, the reduction in amyloid-β_42_ may be a result of a “survivor effect” where in the presence of Dp(16)2Tyb duplication, only mice that had low APP/amyloid-β_42_ may be able to survive to 6-months of age. Further experiments in an alternative model system are required to investigate this finding.

**Fig. 4.**
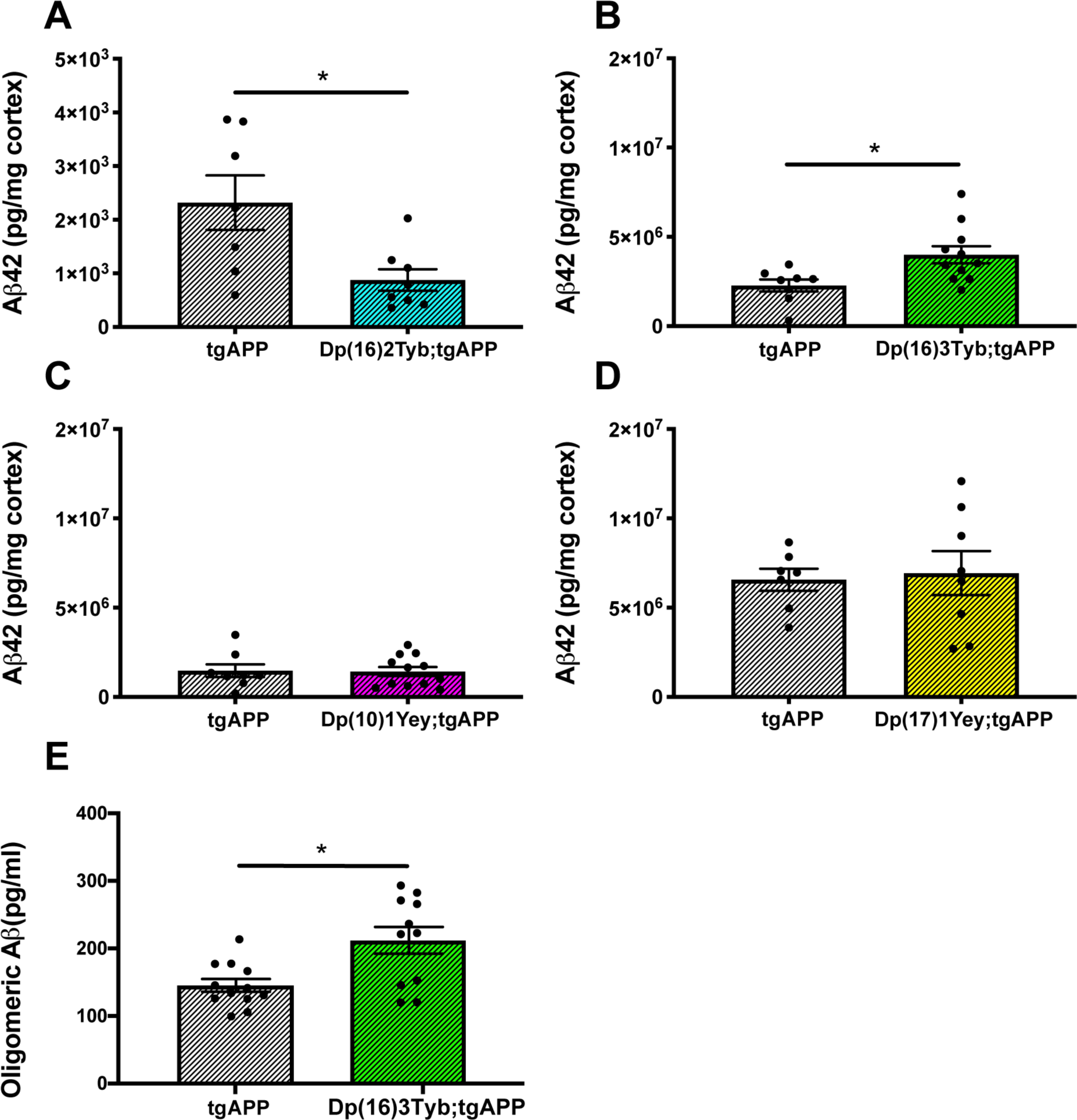
The effect of DS segmental duplication models on insoluble amyloid-β_42_ in the cortex. **(A-D)** Cortical proteins from 6- and 12-month old mice were fractionated and 5 M guanidine hydrochloride, soluble Aβ was quantified by Meso Scale Discovery Assay. Error bars show SEM, data points are independent mice. **(A)** In Dp(16)2Tyb;tgAPP (n = 8, 3 male and 5 female) mice at 6 months of age, median amyloid-β_42_ abundance is significantly decreased (U(N_Dp(16)2Tyb;tgAPP_ = 8, N_tgAPP_ = 7,) = 8, p = 0.021) in cortex compared to tgAPP (n = 7, 2 male and 5 female) littermates. After systematic outlier testing one tgAPP sample was excluded prior to analysis. **(B)** In Dp(16)3Tyb;tgAPP (n = 11, 7 male and 4 female) mice at 12 months of age, amyloid-β_42_ abundance is significantly increased (F(1,13) = 4.656, p = 0.05) compared to tgAPP (n = 8, 5 male and 3 female) littermates. After systematic outlier testing two tgAPP samples were excluded prior to analysis. **(C)** In Dp(10)1Yey;tgAPP (n = 12, 6 male and 6 female) mice at 12 months of age, amyloid-β_42_ abundance (F(1,14) = 0.027, p = 0.872) did not significantly differ compared to tgAPP (n = 8, 5 male and 3 female) littermates. After systematic outlier testing one tgAPP sample and two Dp(10)1Yey;tgAPP samples were excluded prior to analysis. **(D)** In Dp(17)1Yey;tgAPP (n = 8, 3 male and 5 female) mice at 12 months of age, amyloid-β_42_ abundance (F(1,9) = 2.115 p = 0.176) did not significantly differ compared to tgAPP (n = 7, 4 male and 3 female) littermates. After systematic outlier testing two tgAPP samples and one Dp(17)1Yey;tgAPP sample were excluded prior to analysis. **(E)** In Dp(16)3Tyb;tgAPP (n = 11, 6 male and 5 female) mice at 3 months of age, median oligomeric Aβ species were significantly increased U(N_Dp(16)3Tyb;tgAPP_ = 11, N_tgAPP_ = 12,) = 104, p = 0.019) compared to tgAPP (n = 12, 8 male and 4 female) littermates.

Significantly elevated levels of insoluble amyloid-β_42_ were observed in the Dp(16)3Tyb;tgAPP mice at 12-months of age in the cortex, compared to tgAPP littermates (**Fig. 4B**). However, at 6-months of age although a small increase in insoluble amyloid-β_42_ was observed this was not statistically significant (**Fig. S3**). An additional copy of the Dp(10)1Yey or Dp(17)1Yey duplications did not alter insoluble amyloid-β_42_ abundance (**Fig. 4 C, D**).

The increased survival of Dp(16)3Tyb;tgAPP mice compared to tgAPP littermates could contribute to this change in phenotype, because Dp(16)3Tyb;tgAPP mice may be able to tolerate a higher APP/amyloid-β_42_ load than tgAPP littermates, without ill effects. Thus to further assess whether the Dp(16)3Tyb duplication increased amyloid-β aggregation, the abundance of multimeric amyloid-β was measured by ELISA in cortex of Dp(16)3Tyb;tgAPP mice and tgAPP littermates at 3 months of age. This time-point is prior to the timing of significant mortality of tgAPP littermates and thus should control for the increased survival of Dp(16)3Tyb;tgAPP mice. Significantly more multimeric amyloid-β was detected in Dp(16)3Tyb;tgAPP mice than tgAPP littermates (**Fig. 4E**). Thus an additional copy of a gene(s) located between *Mir802* and *Zbtb21* promotes amyloid-β aggregation in the brain in both young and old mice.

An additional copy of Hsa-21 homologues from the Dp(16)2Tyb, Dp(16)3Tyb or Dp(17)1Yey regions did not significantly alter the abundance of insoluble amyloid-β_40_, soluble amyloid-β_40_ or soluble amyloid-β_42_ (**Fig. S4, S5 A, B, D**). Thus the increase in insoluble amyloid-β_42_ abundance caused by the extra copy of the Dp(16)3Tyb duplication likely occurs because of enhanced amyloid-β_42_ aggregation or impaired clearance. An additional copy of the Dp(10)1Yey region led to an increase in Tris soluble amyloid-β_42_ at 12 months of age but this was not seen at 6 months of age (**Fig. S5 C**). The Dp(10)1Yey region did not alter the abundance of insoluble or soluble amyloid-β_40_ at either time-point.

### Additional copies of Hsa21 homologues do not modulate deposition of amyloid-β in the brain

To determine if the changes in amyloid-β_42_ aggregation in the Dp(16)2Tyb-tgAPP and Dp(16)3Tyb-tgAPP models also affected deposition of the peptide we undertook histological studies of brain at 6- and 12-months of age.

A non-significant trend for decreased amyloid-β deposition was observed in the hippocampus at 6- and 12-months of age in the presence of the Dp(16)2Tyb duplication (**Fig. 5A, E**). This small decrease in amyloid-β deposition in the Dp(16)2Tyb;tgAPP mice is consistent with the decrease in aggregated amyloid-β_42_ detected by biochemistry. The high mortality of Dp(16)2Tyb;tgAPP mice meant too few animals survived for a sufficiently powered biochemical study of the solubility/aggregation on amyloid-β. We note that phenotypes in this cross may be the results of a survivor effect.

**Fig. 5.**
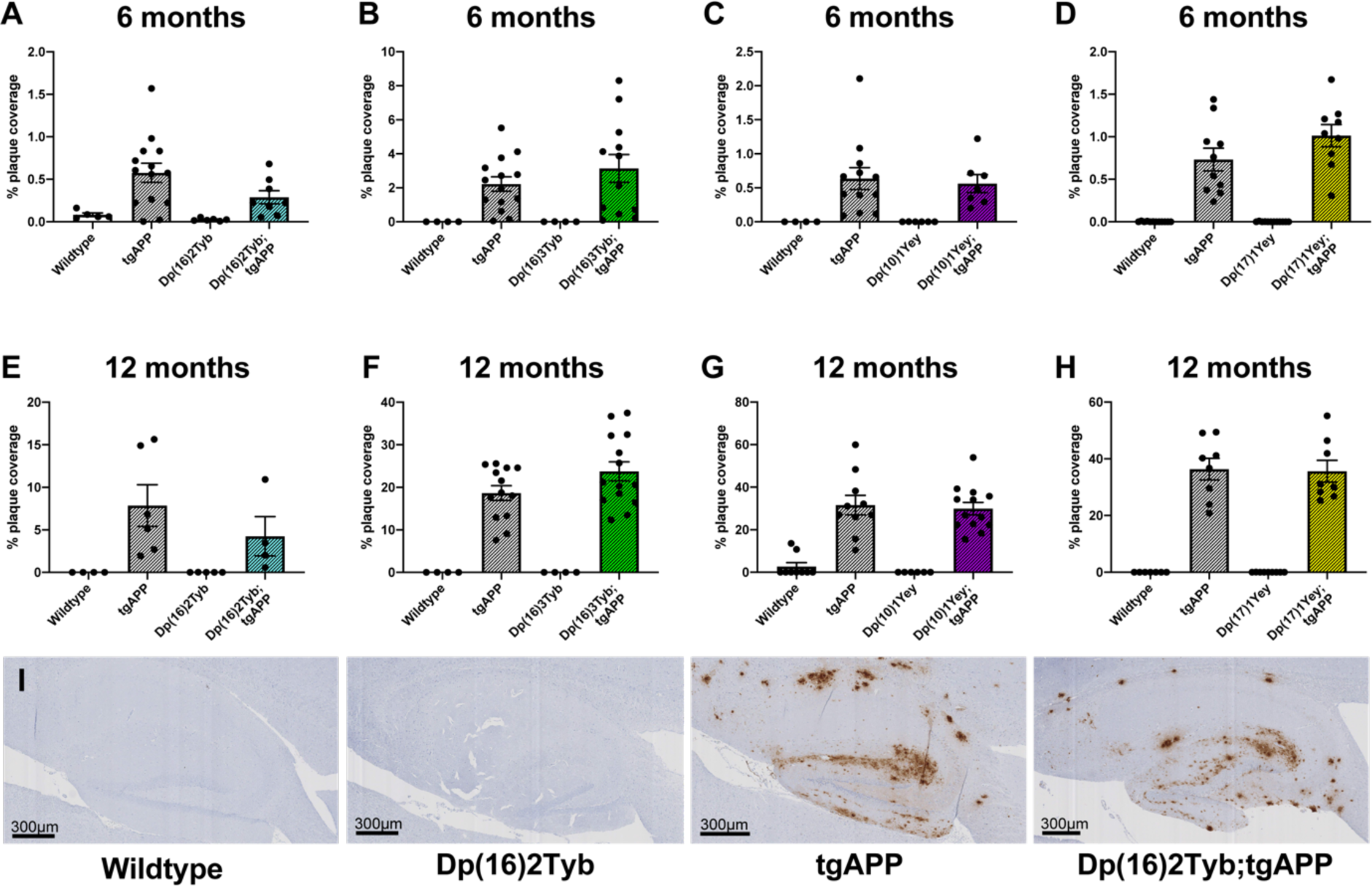
The effect of DS segmental duplication models on amyloid-β in the hippocampus. Amyloid-β deposition in the hippocampus was quantified at (**A-D**) 6- and (**E-H**) 12-months of age in male and female mice, percentage of the region covered by stain was calculated. Error bars show SEM, data points are independent mice. **(A)** No significant difference in amyloid-β deposition in the hippocampus was detected at 6-months of age in Dp(16)2Tyb;tgAPP compared with tgAPP controls (F(1,17) = 2.372, p = 0.142) detected with 82E1 primary antibody. After systematic outlier testing one tgAPP sample was excluded prior to analysis. Dp(16)2Tyb;tgAPP female n=5, male n=3; tgAPP female n=7, male n=7. **(B)** No significant difference in amyloid-β deposition in the hippocampus was detected at 6-months of age in Dp(16)3Tyb;tgAPP compared with tgAPP controls (*U*(N_Dp(16)3Tyb;tgAPP_ = 12, N_tgAPP_ = 14,) = 71, p = 0.5267) detected with 82E1 primary antibody. Dp(16)3Tyb;tgAPP female n=6, male n=6; tgAPP female n=7, male n=7. **(C)** No significant difference in amyloid-β deposition in the hippocampus was detected at 6-months of age in Dp(10)1Yey;tgAPP compared with tgAPP controls (F(1,14) = 0.001, p = 0.972) detected with 4G8 primary antibody. Dp(10)1Yey;tgAPP female n=4, male n=3; tgAPP female n=5, male n=7. **(D)** No significant difference in amyloid-β deposition in the hippocampus was detected at 6-months of age in Dp(17)1Yey;tgAPP compared with tgAPP controls (F(1,14) = 1.785, p = 0.203) detected with 4G8 primary antibody. Dp(17)1Yey;tgAPP female n=5, male n=4; tgAPP female n=6, male n=4. **(E)** No significant difference in amyloid-β deposition in the hippocampus was detected at 12-months of age in Dp(16)2Tyb;tgAPP compared with tgAPP controls (F(1,5) = 1.908, p = 0.226) detected with 82E1 primary antibody. Dp(16)2Tyb;tgAPP female n=3, male n=1; tgAPP female n=3, male n=3. **(F)** No significant difference in amyloid-β deposition in the hippocampus was detected at 12-months of age in Dp(16)3Tyb;tgAPP compared with tgAPP controls (F(1,22) = 2.012, p = 0.170) detected with 82E1 primary antibody. Dp(16)3Tyb;tgAPP female n=8, male n=6; tgAPP female n=7, male n=7. **(G)** No significant difference in amyloid-β deposition in the hippocampus was detected at 12-months of age in Dp(10)1Yey;tgAPP compared with tgAPP controls (F(1,18) = 0.131, p = 0.722) detected with 4G8 primary antibody. Dp(10)1Yey;tgAPP female n=7, male n=6; tgAPP female n=6, male n=4. **(H)** No significant difference in amyloid-β deposition in the hippocampus was detected at 12-months of age in Dp(17)1Yey;tgAPP compared with tgAPP controls (F(1,11) = 0.021, p = 0.886) detected with 4G8 primary antibody. Dp(17)1Yey;tgAPP female n=5, male n=3; tgAPP female n=5, male n=3. **(I)** Representative image of Dp(16)2Tyb;tgAPP hippocampus at 12-months of age.

No change in amyloid-β deposition in the hippocampus was observed in Dp(16)3Tyb;tgAPP, Dp(10)1Yey;tgAPP and Dp(17)1Yey;tgAPP mice compared with tgAPP littermates at either time-point (**Fig. 5 B, C, D, F, G, H**). Similarly, no significant changes in deposition of amyloid-β in the cortex was observed at 6-months or 12-months of age in the Dp(16)3Tyb;tgAPP, Dp(10)1Yey;tgAPP or Dp(17)1Yey;tgAPP compared to tgAPP littermates (**Fig. S6**). Thus increased amyloid-β_42_ aggregation caused by Dp(16)3Tyb does not result in a robust increase in deposition of amyloid within the brain and the Dp(10)1Yey and Dp(17)1Yey duplications are not sufficient to modulate amyloid-β accumulation.

### The rescue of APP transgene-associated mortality by segmental duplication of *Prmt2* and *Pdxk* is not caused by a suppression of seizures

Elevated mortality in *APP* transgenic mice such as the J20 animals studied here is likely caused by the occurrence of seizures and raised subclinical seizure activity in this overexpression model. The *Prmt2* and *Pdxk* region which is duplicated in the Dp(10)1Yey mouse model of DS carries *Cstb*, which encodes the enzyme cystatin B. Loss of function of this gene causes elevated seizure activity (*20*). Thus an additional copy of this gene may decrease the occurrence of tgAPP-associated seizures in the Dp(10)1Yey model and this may underlie the rescue of mortality in Dp(10)1Yey;tgAPP mice. To test this hypothesis, we measured the occurrence of seizures in tgAPP and Dp(10)1Yey;tgAPP male mice at 3-4 months of age. The frequency and duration of seizures associated with the *APP* transgene was not altered by the Dp(10)1Yey region (**Fig. 6**). Thus the rescue of mortality in Dp(10)1Yey;tgAPP double mutant mice is downstream or independent of seizure occurrence. This suggests that 3 copies of *Cstb* do not modify tgAPP seizure occurrence, consistent with a previous report that showed an increase in *Cstb* copy number does not alter picrotoxin-induced seizure thresholds (*21*).

**Fig. 6.**
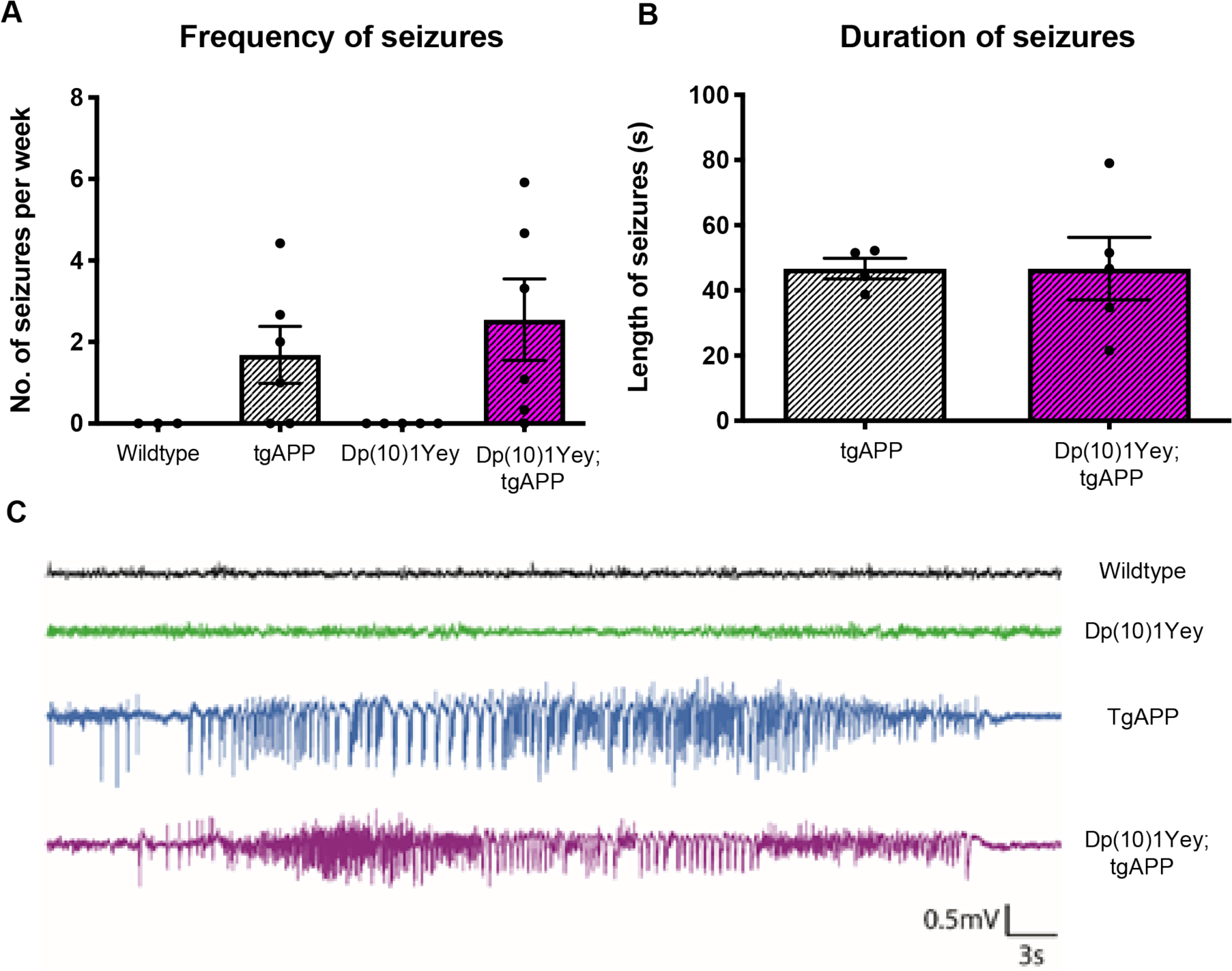
The Dp(10)1Yey segmental duplication did not alter the frequency or duration of APP transgene-associated seizures. **(A)** The frequency and **(B)** duration of seizure-like events was measured in male mice by cortical EEG using implanted electrodes in freely moving mice in the home cage. **(A)** The frequency (T-test p = 0.60, tgAPP n = 6, Dp(10)1Yey;tgAPP n = 6 all male) and **(B)** duration (T-test p = 0.64, tgAPP n = 6, Dp(10)1Yey;tgAPP n = 6) of seizure-like events was not altered by the duplication of the Dp(10)1Yey region. **(C)** Example EEG trace of a seizure-like event in tgAPP and Dp10;tgAPP animals with control time-matched wildtype and Dp(10)1Yey littermates traces for comparison. Error bars show SEM, data points are independent mice.

## DISCUSSION

People who have DS are highly prone to develop early-onset dementia caused by AD (*2*). Here we have used mouse models of DS and tgAPP mouse models of amyloid-β deposition to understand how sub-regions of Hsa21 modulate AD-related biology. Our data suggest that gene(s) from multiple regions of the chromosome affect APP/amyloid-β associated mortality, amyloid-β aggregation and deposition, independently of changes in FL-APP abundance. Moreover, we show that changes in mortality occur independently of an alteration in seizure duration or frequency in the Dp(10)1Yey segmental duplication model, despite this model having 3 copies of epilepsy related gene *Cstb*.

Mouse models are tools to understand the human condition, and within all model systems lie limitations. In this study we have modelled human trisomy by duplication of mouse orthologues. Thus, human genes/isoforms without a mouse equivalent have not been studied. We have determined how an additional copy of sub-groups of mouse Hsa21 orthologues affect biology but we have not determined how these groups of genes may interact with each other. Thus, some of the gene-interactions that occur in people who have DS are not modelled here. Also, we have not studied the effect of an extra copy of the 45 genes on Hsa21 located near to *App* on APP/amyloid-β biology due to the potential confounding effect of increased copy number of endogenous mouse *App*. Additionally, we used an over-expressing APP transgenic model which has high/non-physiological levels of APP and its cleavage fragments, thus modifying effects of some Hsa21-dosage sensitive genes may be masked.

In our previous study crossing an alternative mouse model of DS (Tc1 mouse) with the J20 tgAPP mouse we found a robust increase in both the aggregation and deposition of amyloid-β in the cortex and hippocampus (*12*). In this study we identified a subregion of Hsa21 that is sufficient to cause an increase in the aggregation but not in deposition of amyloid-β. Additionally this increase was not as large as observed in the Tc1;tgAPP mice. Our data suggest that either a human specific gene/isoform, a gene(s) located close to *App*, or a combination of multiple genes across the chromosome is required to robustly increase both aggregation and deposition of amyloid-β reported in our previous model system. Some DS-associated phenotypes result from the combined effect of an extra copy of multiple genes on chromosome 21 (*13, 22*). This may occur via the effect of mass-action of an extra copy of many genes, perhaps via a global impairment in proteostasis as has been recently suggested to occur in DS (*23*). Further work is required to investigate the effect of this mechanism and how an extra copy of the genes located near to *App* effect APP/amyloid-β biology.

Moreover, we observe a significant decrease in aggregation, and non-significant trend for decreased deposition, of amyloid-β caused by an additional copy of the Dp(16)2Tyb region. We note the Tc1 mouse model does not have an additional copy of the genes duplicated in the Dp(16)2Tyb model, which likely underlies some of the phenotypic differences between these two inter-crosses. All phenotypes observed in the Dp(16)2Tyb cross may be the result of survivor effect because of the highly elevated mortality of the Dp(16)2Tyb;tgAPP mice. Further work in an alternative model system is required to investigate how the Dp(16)2Tyb region modulates aggregation and deposition of amyloid-β.

Mortality in APP transgenic mouse models including the J20 line is caused by seizures (*16*), likely by a process similar to Sudden Unexpected Death in Epilepsy (SUDEP). SUDEP is thought to occur because of a post-seizure response which slows electrical activity in the brain-stem leading to a fatal suppression of respiratory and cardiac output (*24, 25*). Here we show that tgAPP associated mortality is rescued by an extra copy of multiple regions of Hsa21, and in Dp(10)1Yey mice this occurs despite the frequent occurrence of seizure activity. This suggests that an increase in copy number of a gene(s) in this region of Hsa21 may protect against SUDEP. Similarly, copy number of genes in the Dp(16)2Tyb region may worsen the incidence, duration or adverse outcome of seizures. Further research will determine the mechanism of action of this region and whether genes in this region contribute to the increased incidence of epilepsy and dementia-associated seizures in people who have DS.

In conclusion, an additional copy of genes on Hsa21 other than *App* modulate APP/amyloid-β biology, including tgAPP associated mortality and amyloid-β aggregation in a transgenic mouse model. Thus, genes on Hsa21 other than *APP* may influence the development and progression of AD in people who have DS, and AD therapies for this important group of individuals must be carefully selected to take this into account. Lastly, our data is consistent with DS-associated phenotypes being the result of human-specific gene effects or the interaction of an extra copy of multiple genes on Hsa21.

## MATERIALS AND METHODS

### Animal welfare and husbandry

Mice were housed in individually ventilated cages (Techniplast) with grade 5, autoclaved dust-free wood bedding, paper bedding and a translucent red “mouse house”. Free-access to food and water was provided. The animal facility was maintained at a constant temperature of 19-23°C with 55 ± 10% humidity in a 12 h light/dark cycle. Pups were weaned at 21 days and moved to standardised same-sex group housing with a maximum of 5 mice per cage.

The following mouse strains were used in this paper, here we show abbreviated name and then the official name and unique Mouse Genome Informatics (MGI) identifier: Dp(16)2Tyb (Dp(16Mis18a-Runx1)2TybEmcf, MGI:5703800), Dp(16)3Tyb (Dp(16Mir802-Zbtb21)3TybEmcf, MGI:5703802), Dp(10)1Yey (Dp(10Prmt2-Pdxk)2Yey, MGI:4461400) and Dp(17)1Yey (Dp(17Abcg1-Rrp1b)3Yey, MGI:4461398).

Mice were maintained by backcrossing males and females to C57BL/6J mice. tgAPP mice (B6.Cg-Tg(PDGFB-APPSwInd)20Lms, MGI:3057148) were maintained by mating tgAPP female mice to C57BL/6J male mice. Experimental cohorts were generated by crossing male or female mice carrying Hsa21 orthologous duplications with male or female tgAPP mice.

Animals were euthanized by exposure to rising carbon dioxide, followed by confirmation of death by dislocation of the neck in accordance with the Animals (Scientific Procedures) Act 1986 (United Kingdom).

### Tissue preparation and western blotting

For analysis of protein abundance in hippocampus and cortex, tissue was dissected under ice cold PBS before snap freezing. Samples were then homogenised in RIPA Buffer (150 mM sodium chloride, 50 mM Tris, 1 % NP-40, 0.5 % sodium deoxycholate, 0.1 % sodium dodecyl sulfate) plus complete protease inhibitors (Calbiochem) by mechanical disruption. Total protein content was determined by Bradford assay or Pierce™ 660nm assay (ThermoFisher). Samples from individual animals were run separately and were not pooled.

Equal amounts of total brain proteins were then denatured in LDS denaturing buffer (Invitrogen) and β-mercaptoethanol, prior to separation by SDS-PAGE gel electrophoresis using precast 4-12 % Bis-Tris gels (Invitrogen). Proteins were transferred to nitrocellulose or PVDF membranes prior to blocking in 5 % milk/PBST (0.05 % Tween-20) or 5-10 % bovine serum albumin (BSA)/PBST. Primary antibodies were diluted in 1 % BSA/PBST, HRP-conjugated secondary anti-rabbit, anti-mouse and anti-goat antibodies (Dako) were diluted 1:10,000 in 1% BSA/PBST. Linearity of antibody binding was confirmed by a 2-fold dilution series of cortical protein samples. Band density was analysed using Image J. Relative signal of the antibody of interest compared to the internal loading control was then calculated, and relative signal was then normalized to mean relative signal of control samples run on the same gel. Mean of technical replicates were calculated and used for ANOVA, such that biological replicates were used as the experimental unit.

Primary antibodies against C-terminal APP (Sigma A8717, 1:10,000), β-actin (Sigma A5441, 1:60,000), and GAPDH (Sigma G9545, 1:200,000), were used.

### Biochemical fractionation of mouse brain tissues

Cortical proteins were fractionated as described in Shankar *et al*. (2009). A half cortex was weighed on a microscale and homogenised in 4 volumes of ice-cold Tris-buffered saline (TBS) (50mM Tris-HCl pH 8.0) containing a cocktail of protease and phosphatase inhibitors (Calbiochem) using a handheld mechanical homogeniser and disposable pestles (Anachem). Samples were then transferred to 1.5ml microfuge tubes (Beckman Coulter #357448), balanced by adding more TBS and centrifuged at 175,000 × g with a RC-M120EX ultracentrifuge (Sorvall) fitted with rotor S100AT5 at 4 °C for 30 mins. Supernatant (the Tris-soluble fraction) was removed and stored at -80 °C. The remaining pellet was homogenised in 5 volumes of ice-cold 1 % Triton-X (Sigma-Aldrich) in TBS (50mM Tris-HCl pH 8.0), balanced and centrifuged at 175,000g for 30 mins at 4 °C. The resultant supernatant (the Triton-soluble fraction) was removed and stored at -80 °C. The pellet was then re-suspended in 8 volumes (by original cortical weight) in TBS (50mM Tris-HCl pH 8.0), containing 5M Guanidine HCl and left overnight at 4 °C on a rocker to ensure full re-suspension, and subsequently stored at -80 °C. A 660 nm protein assay (ThermoFisher) was performed to determine protein concentration for normalisation following ELISA assay. For 3-month old animals, hippocampal TBS-soluble protein fractions were prepared according to Hölttä et al. (2013). Total mouse hippocampus from the left hemisphere was homogenised using a mechanical homogeniser and disposable pestles (Anachem) in 100μl TBS (50mM Tris-HCl pH 8.0) containing protease and phosphatase inhibitors (Calbiochem). The homogenate was then centrifuged at 4 °C for 30 mins at 16,000 × g. The supernatant (Tris-soluble fraction) was removed and aliquoted to store at -80 °C. 660 nm protein assay was used to assess protein concentration.

### Quantification of Aβ production by Meso Scale Discovery Assay

Aβ38, Aβ40 and Aβ42 product levels were quantified on Multi-Spot 96 well plates pre-coated with anti-Aβ38, Aβ40, and Aβ42 antibodies obtained from Janssen Pharmaceutica using multiplex MSD technology, as described before in (*27*).

### Sandwich ELISA for Aβ oligomers

Oligomeric Aβ was detected using a sandwich ELISA adapted from (*28*) using 82E1 monoclonal antibody for both capture and detection to exclude monomeric Aβ from detection. Briefly, total hippocampus from 3-month old animals was homogenized using a mechanical homogenizer and disposable pestles (Anachem) in approximately 5 volumes of ice-cold Tris-buffered saline (TBS) (50 mM Tris-HCL, pH 8.0) containing a cocktail of protease and phosphatase inhibitors (Calbiochem). Homogenates were centrifuged at 16,000 x g at 4 °C for 30 min, the resultant supernatant (the soluble TBS fraction) was stored at – 80 °C. For the sandwich ELISA, samples were incubated on 1μg/ml 96 well plate-bound 82E1 capture antibody followed by incubation with 0.75μg/ml Biotinylated 82E1 detection antibody. The plate was washed 5 times with PBS-T between each step. The standard curve was made up of 2-fold dilutions, ranging from 3678 to 3.6 pg/ml of a synthetic dimer with the Aβ1-11 sequence (DAEFRHDSGYE) C-terminally linked via a cysteine residue (*28*) was ordered from rPeptide and supplied by Stratech.

NeutrAvidin-HRP (ThermoFisher Scientific) was added to the plate after which a colour reaction was generated by incubation with TMB (3,3’,5,5’-tetramethylbenzidine) colorimetric substrate (1-Step Ultra TMB-ELISA Substrate Solution, ThermoFisher Scientific). The reaction was stopped with 2 M H_2_SO_4_ and the plate read at 405nm with a Tecen Infinite M1000 plate-reader. The average reading from the three technical replicates and the standard deviation for each standard were calculated to determine each one’s percent coefficient of variation (%CV). The lower limit of detection (LLOD) was calculated as 2.5 times the standard deviation of the blank standard. The acceptable limits for each sample’s %CV value were then taken to fall between the LLOD percentage and 10%. The standard curve was plotted and the region of linearity with the best R-square value was determined empirically. The estimated concentration for each standard was calculated from the plot equation and was divided by the assigned concentration to assess their similarity (% backfit). The lower limit and upper limits of quantification (LLOQ and ULOQ) were set to fall between 80 and 120% of the % backfit values. The concentration of each sample was then determined according to the above limits.

### Implantation of subcutaneous EEG transmitters

Animals were anesthetized using 4% 1mg/l isoflurane vapour (Isothesia, Henry Schein Animal Health, UK), then maintained on approximately 1.5% isoflurane vapour. All animals received saline bolus prior to and after surgery, and Metacam (Boehringer) and buprenorphine (Vibec) analgesia. Intracranial electrodes were implanted 1 mm into the right parietal cortex (−2.06, +2.50, with reference to bremga) and right motor cortex (+1.00, +1.5, with reference to bremga), via holes drilled through the skull. Electrodes were connected to single channel radio-transmitter (A3028B-AA, frequency dynamic range of 0.3-160 Hz, sampling rate of 512 per second (SPS) 600 hours recording or A3028C-AA frequency dynamic range of 0.3-80 Hz, 256 SPS, 950 hours recording) and battery implanted subcutaneously in the animals body (Open Source Instruments), such that EEG recordings could be made continuously from an untethered animal in its home-cage. EEG was recorded in freely moving animals continuously over a period of 6-21 days inside a faraday cage, signals recorded using LWDAQ software, in NDF file format.

### Automated seizure and sub-clinical epileptiform activity detection

Automated seizure detection using an ECP16V1 processing script Neuroarchiver v101 (software available at http://alignment.hep.brandeis.edu/Software/), was used which calculates numerical metrics for 6 EEG properties: power, coastline, intermittency, coherence, asymmetry, and rhythm, over each second of EEG (source code available at: http://www.opensourceinstruments.com/Electronics/A3018/Seizure_Detection.html). A library of 1-second EEG segments containing visually identified examples of seizures, sub-clinical epileptiform activity (SCEA), baseline EEG and movement artefacts was created using the ‘event classifier’ feature in Neuroarchiver v101. The threshold used was 0.04 for SCEA and 0.1 for seizures. All events were then checked manually to exclude false positives. Seizures were defined as high frequency and high amplitude spiking activity lasting longer than 10s. Any spiking lasting less than 10 seconds was classified as SCEA. Event frequency was normalised by the number of days of recording.

### Immunohistochemistry of mouse brain

Half brains were immersion fixed in 10 % buffered formal saline (Pioneer Research Chemicals) for a minimum of 48 hours prior to being processed to wax (Leica ASP300S tissue processor). The blocks were trimmed laterally from the midline by ∼0.9-1.4 mm to give a sagittal section of the hippocampal formation. Two 4 μm sections 40 μm apart were analysed. The sections were pretreated with 98 % formic acid for 8 minutes, followed by washing. The slides were wet loaded onto a Ventana XT for staining (Ventana Medical Systems, Tuscon, AZ, USA). The protocol included the following steps: heat induced epitope retrieval (mCC1) for 30 minutes in Tris Boric acid EDTA buffer (pH 9.0), superblock (8mins) and manual application of 100μl of directly biotinylated mouse monoclonal IgG1 antibodies against Aβ (82E1, IBL, 0.2 μg/ml or 4G8, Millipore, 2 μg/ml) for 8 hours. The staining was visualised using the Ventana DabMap kit (iView DAB, Ventana Medical Systems), followed by 4mins of haematoxylin and blueing. Alternatively, for staining of Beta-amyloid, slides were incubated with mouse monoclonal 6F/3D (Dako 1:50) followed by Iview Ig secondary antibody (Ventana Medical Systems). The sections were dehydrated, cleared and mounted in DPX prior to scanning (Leica SCN400F scanner). All images were analysed using Definiens Tissue Studio software (Definiens Inc). 6F/3D stained slides were photographed (ImageView II 3.5 Mpix digital camera) and composed with Adobe Photoshop so that the entire cortex could be analysed. The same thresholds for staining intensity were then used to quantify the area covered by DAB stain using Volocity image analysis software (Perkin Elmer).

### Statistical analysis

Data were analysed as indicated in figure legends by either two-tailed students T-test (single variable study), univariate ANOVA (to control for multiple variables) or by Mann-Whitney U, a non-parametric test, in cases where sample groups failed a Levene’s test for equality of distribution between samples. For ANOVA, between-subject factors were trisomy and sex, with age in days included as a covariate. For cases when the number of technical replicates varied between subjects, subject means were calculated and used in the ANOVA. For MSD assays, fractionation batch was included as a covariate. For MSD assays and Aβ immunohistochemistry data, data points which were greater than three times the interquartile range of its group were excluded from analysis and reported in the figure legend.

## Acknowledgement

F.K.W. is supported by the UK Dementia Research Institute which receives its funding from DRI Ltd, funded by the UK Medical Research Council, Alzheimer’s Society and Alzheimer’s Research UK and by an Alzheimer’s Research UK Senior Research Fellowship. FKW also received funding that contributed to the work in this paper from Epilepsy Research UK and the MRC via CoEN award MR/S005145/1. J.L.T. was funded by an Alzheimer’s Society PhD studentship awarded to F.K.W. and EMCF. L.J.P. was funded by an Alzheimer’s Research UK PhD studentship awarded to F.K.W. and E.M.C.F. The authors were funded by a Wellcome Trust Strategic Award (grant number: 098330/Z/12/Z) awarded to The London Down Syndrome (LonDownS) Consortium (V.L.J.T., and E.M.C.F). Additionally, the authors were funded by a Wellcome Trust Joint Senior Investigators Award (V.L.J.T. and E.M.C.F., grant numbers: 098328, 098327), the Medical Research Council (programme number U117527252; awarded to V.L.J.T). V.L.J.T. is also funded by the Francis Crick Institute which receives its core funding from the Medical Research Council (FC001194), Cancer Research UK (FC001194) and the Wellcome Trust (FC001194). R.C.W. holds a Senior Research fellowship funded by the Worshipful Company of Pewterers and an Epilepsy Research UK Fellowship (F1401).

We thank Professor S. Schorge (Royal Society fellowship URF (UF140596)), Dr. Amanda Heslegrave and Dr. Matthew Ellis for assistance with this project.

F.K.W. has undertaken consultancy for Elkington and Fife Patent Lawyers unrelated to the work in the manuscript.

J.T undertook biochemical and histological experiments, undertook data analysis and wrote the manuscript. P.M. undertook data analysis and wrote the manuscript. H.T.W, E.R. and L.J.P. undertook biochemical and histological experiments, S.N undertook histological experiments and K.C. undertook biochemical experiments and genotyping. R.C.W, S.S and M.C.W. oversaw the EEG study and R.C.W and L.J.P undertook the EEG experiments and data analysis. E.M.C.F. and V.L.J.T designed and supervised the study and wrote the manuscript. F.K.W. designed and supervised the study, undertook data analysis and wrote the manuscript.

## Supplementary material

**Fig. S1.**
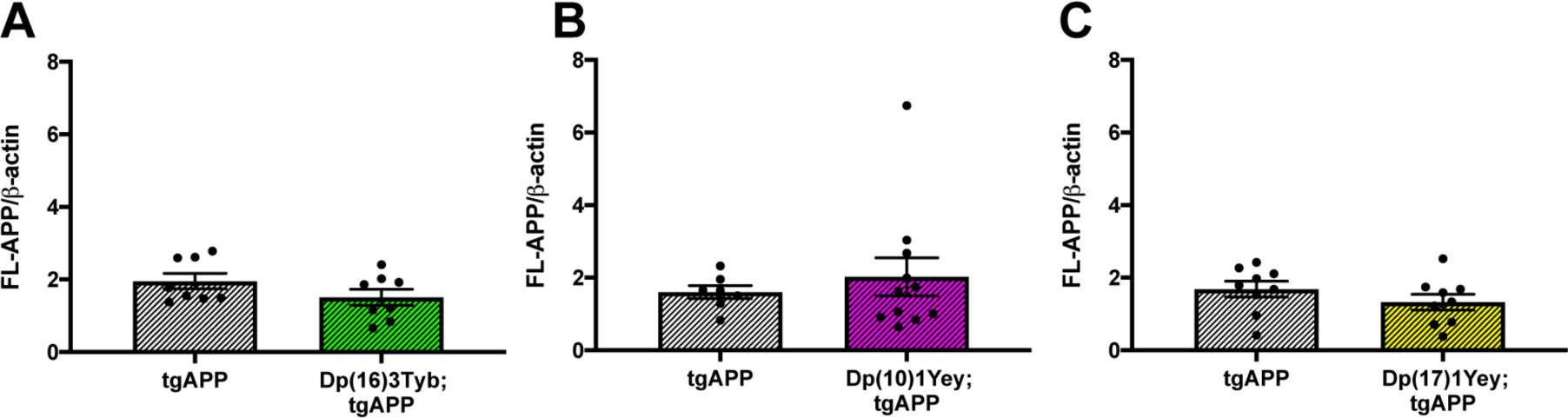
Abundance of FL-APP is not affected by duplications in the DS mouse models at 3-months of age in the cortex. The abundance of full-length APP (FL-APP) relative to β-actin loading control was measured by western blot using A8717 primary antibody in the cortex at 3-months of age in male and female mice. **(A)** There was no difference in FL-APP level between Dp(16)3Tyb;tgAPP (n = 8, 4 male and 4 female) and tgAPP (n = 8, 4 male and 4 female) littermate controls (F(1,12) = 1.896, p = 0.194). **(B)** No difference in FL-APP level between Dp(10)1Yey;tgAPP (n = 11, 7 male and 4 female) and tgAPP (n = 7, 3 male and 4 female) littermate controls (F(1,14) = 0.520, p = 0.576). **(C)** No difference in FL-APP level between Dp(17)1Yey;tgAPP mice (n = 9, 6 male and 3 female) and tgAPP littermate controls (F(1,14) = 0.500, p = 0.491). Error bars show SEM, data points are independent mice.

**Fig. S2.**
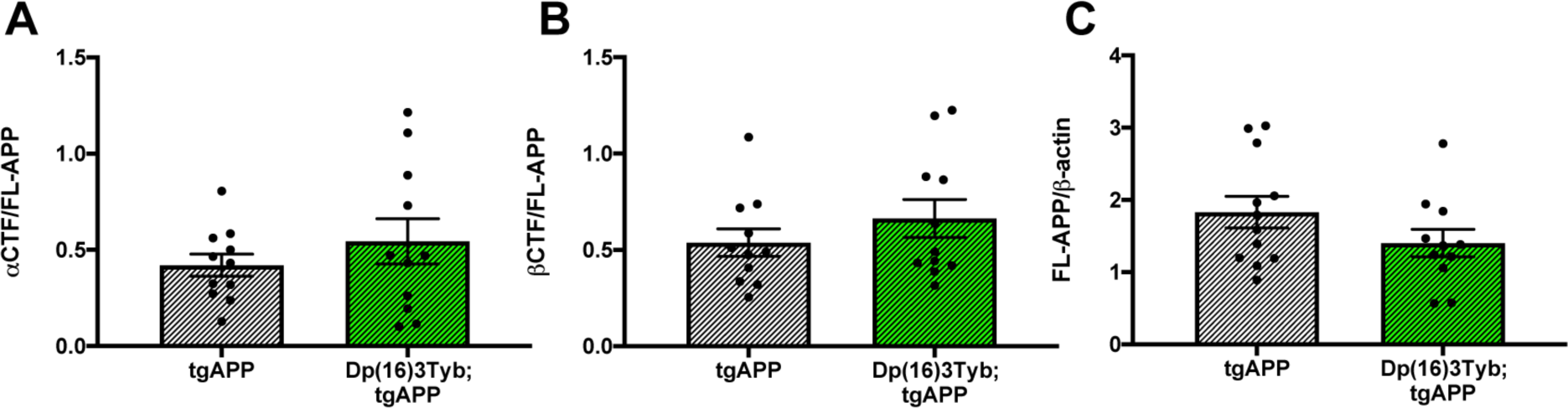
Abundance of FL-APP and CTFs is not altered by the Dp(16)3Tyb duplication at 3-months of age in the hippocampus. The abundance of full-length APP (FL-APP) relative to β-actin loading control, and APP β-C-terminal fragment (β-CTF) and APP α-C-terminal fragment (α-CTF) relative to full-length APP (FL-APP) was measured by western blot using A8717 primary antibody in the hippocampus at 3-months of age in female and male mice. No difference in, or **(A)** α-CTF (F(1,10) = 0.019, p = 0.892) **(B)** β-CTF (F(1,11)= 1.493, p = 0.247) **(C)** FL-APP (F(1,11) = 1.305, p = 0.277) abundance between Dp(16)3Tyb;tgAPP (n = 12, male = 8 and female = 4) and tgAPP (n = 12, male = 8 and female = 4) littermate controls. Error bars show SEM, data points are independent mice.

**Fig. S3.**
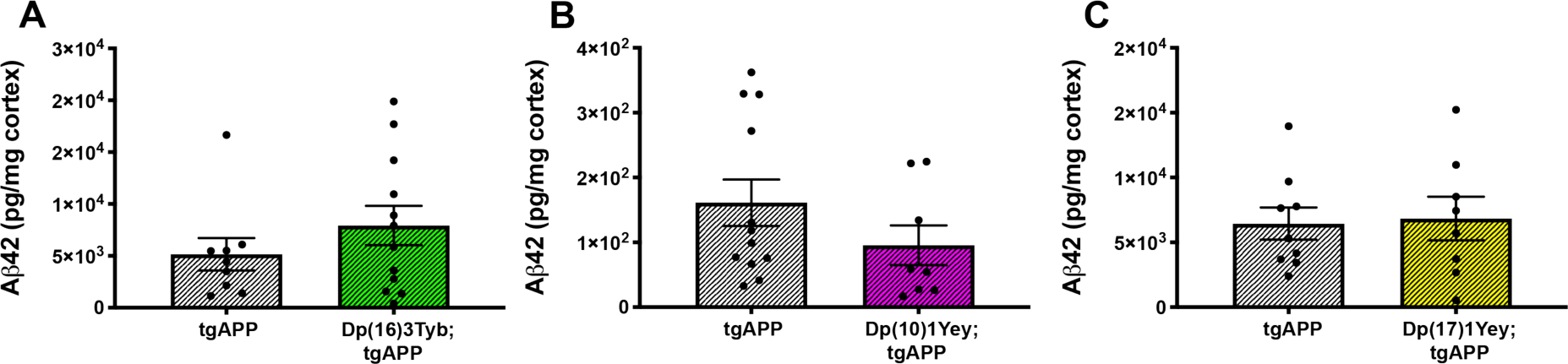
An additional copy of Hsa-21 homologues from the Dp(16)3Tyb, Dp(10)1Yey, or Dp(17)1Yey regions did not alter the abundance of insoluble amyloid-β_42_ in cortex at 6-months of age. **(A)** In Dp(16)3Tyb;tgAPP mice (n = 12, 7 male and 5 female) insoluble amyloid-β_42_ abundance (F(1,16) = 1.851 p = 0.192) did not significantly differ from tgAPP (n = 10, 3 male and 7 female) littermates at 6-months of age. **(B)** In Dp(10)1Yey;tgAPP (n = 8, 3 male and 5 female) mice insoluble amyloid-β_42_ abundance (F(1,14) = 2.990 p = 0.106) did not significantly differ from tgAPP (n = 12, 7 male and 5 female) littermates at 6-months of age. **(C)** In Dp(17)1Yey;tgAPP (n = 8, 4 male and 4 female) mice insoluble amyloid-β_42_ abundance (F(1,11) = 0.299 p = 0.596) did not significantly differ from tgAPP (n = 9, 4 male and 5 female) littermates at 6-months of age. Error bars show SEM, data points are independent mice.

**Fig. S4.**
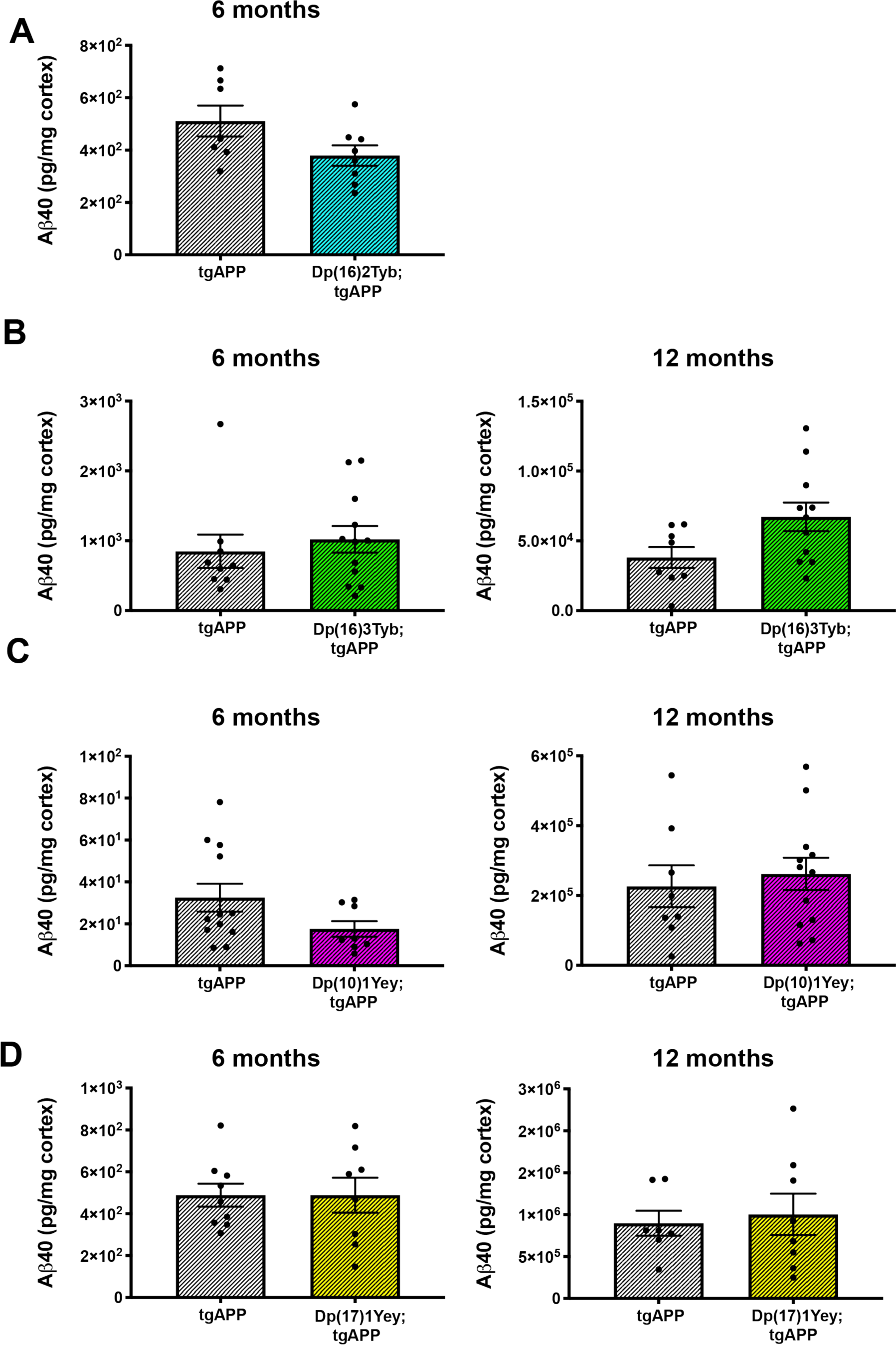
An additional copy of Hsa-21 homologues from the Dp(16)2Tyb, Dp(16)3Tyb, Dp(10)1Yey, or Dp(17)1Yey regions did not alter the abundance of insoluble amyloid-β_40_ in cortex at 6- or 12- months of age. **(A)** In Dp(16)2Tyb;tgAPP (n = 8, 3 male and 5 female) mice insoluble amyloid-β_40_ abundance (F(1,9) = 0.3.739 p = 0.085) did not significantly differ from tgAPP (n = 7, 2 male and 5 female) littermates at 6-months of age. **(B)** In Dp(16)3Tyb;tgAPP (n = 12, 7 male and 5 female) mice insoluble amyloid-β_40_ abundance (F(1,16) = 0.654 p = 0.431) did not significantly differ from tgAPP (n = 10, 3 male and 7 female) littermates at 6-months of age. In Dp(16)3Tyb;tgAPP mice (n = 11, 7 male and 4 female) insoluble amyloid-β_40_ abundance (F(1,13) = 2.776, p = 0.120) did not significantly differ from tgAPP (n = 8, 5 male and 3 female) littermates at 12-months of age. **(C)** In Dp(10)1Yey;tgAPP (n = 8, 3 male and 5 female) mice insoluble amyloid-β_40_ abundance (F(1,14) = 3.417 p = 0.086) did not significantly differ from tgAPP (n = 12, 7 male and 5 female) littermates at 6-months of age. In Dp(10)1Yey;tgAPP (n = 12, 6 male and 6 female) mice insoluble amyloid-β_40_ abundance (F(1,14) = 1.112, p = 0.307) did not significantly differ from tgAPP (n = 8, 5 male and 3 female) littermates at 12-months of age. **D)** In Dp(17)1Yey;tgAPP (n = 8, 4 male and 4 female) mice insoluble amyloid-β_40_ abundance (F(1,11) = 0.498, p = 0.495) did not significantly differ from tgAPP (n = 9, 4 male and 5 female) littermates at 6-months of age. In Dp(17)1Yey;tgAPP (n = 8, 3 male and 5 female) mice insoluble amyloid-β_40_ abundance (F(1,9) = 0.645, p = 0.443) did not significantly differ from tgAPP (n = 7, 4 male and 3 female) littermates at 12-months of age. Error bars show SEM, data points are independent mice.

**Fig. S5.**
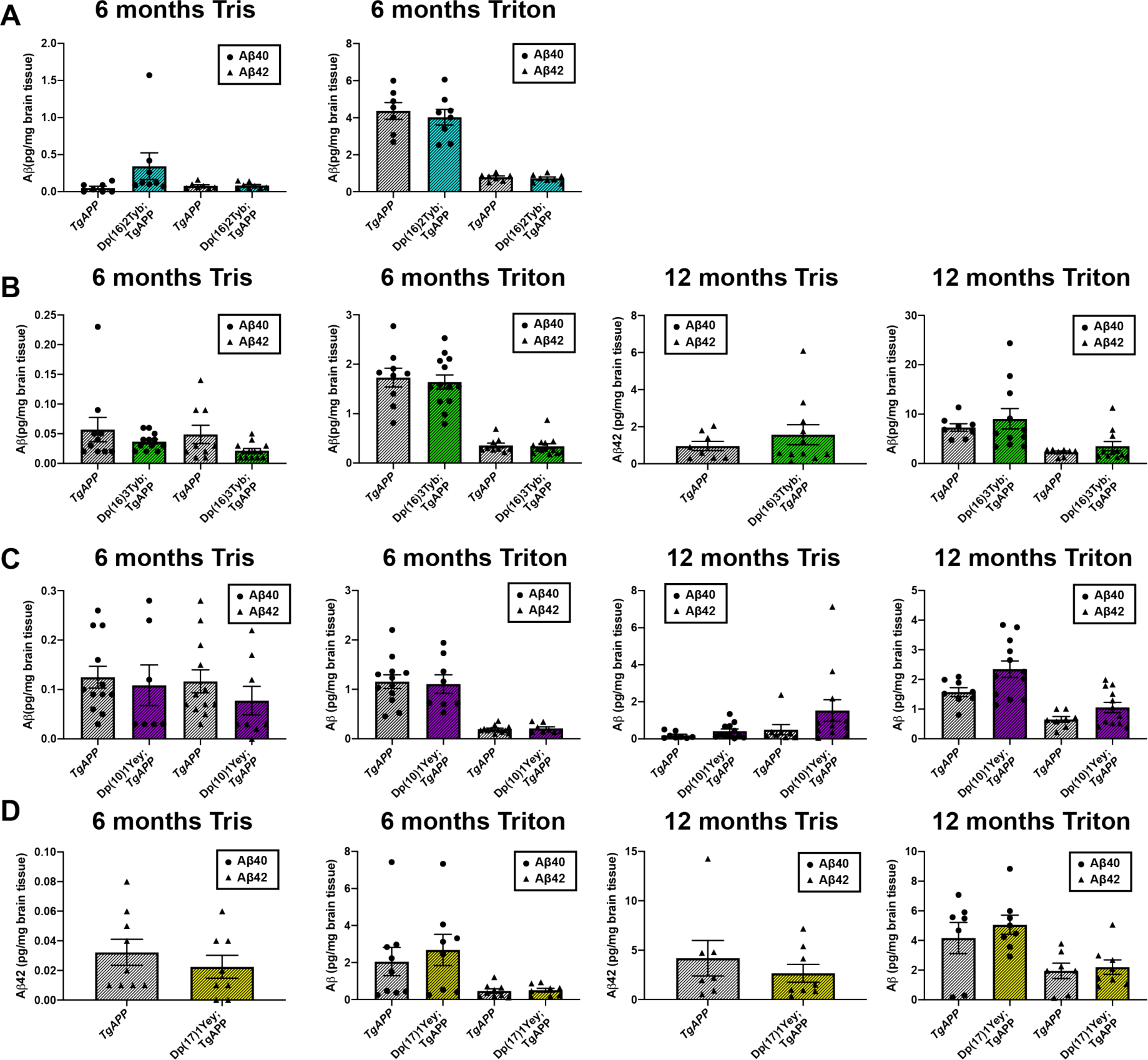
The effect of an additional copy of Hsa-21 homologues from the Dp(16)2Tyb, Dp(16)3Tyb, Dp(10)1Yey, or Dp(17)1Yey regions on soluble Tris and Triton amyloid-β_40_ and amyloid-β_42_ in the cortex at 6- or 12- months of age. **(A)** In Dp(16)2Tyb;tgAPP mice in the soluble Tris fraction, amyloid-β_4o_ abundance (F(1,9) = 1.131, p = 0.315) and amyloid-β_42_ abundance (F(1,9) = 0.261, p = 0.622) did not significantly differ from tgAPP littermates at 6-months of age. In Dp(16)2Tyb;tgAPP mice in the soluble Triton fraction, amyloid-β_4o_ abundance (F(1,9) = 0.08, p = 0.784) and amyloid-β_42_ abundance (F(1,9) = 0.224, p = 0.647) did not significantly differ from tgAPP littermates at 6-months of age. Dp(16)2Tyb;tgAPP (n = 8, 3 male and 5 female) tgAPP (n = 7, 2 male and 5 female). **(B)** In Dp(16)3Tyb;tgAPP mice in the soluble Tris fraction, amyloid-β_4o_ abundance (F(1,16) = 0.789, p = 0.387) and amyloid-β_42_ abundance (U(N_Dp(16)3Tyb;tgAPP_ = 12, N_tgAPP_ = 10,) = 44, p = 0.341) did not significantly differ from tgAPP littermates at 6-months of age. In Dp(16)3Tyb;tgAPP mice in the soluble Triton fraction, amyloid-β_4o_ abundance (F(1,16) = 0.006, p = 0.940) and amyloid-β_42_ abundance (F(1,16) = 0.006, p = 0.844) did not significantly differ from tgAPP littermates at 6-months of age Dp(16)3Tyb;tgAPP (n = 12, 7 male and 5 female) tgAPP (n = 10, 3 male and 7 female). In Dp(16)3Tyb;tgAPP mice in the soluble Tris fraction, amyloid-β_42_ abundance (F(1,13) = 0.332, p = 0.574) did not significantly differ from tgAPP littermates at 12-months of age. Amyloid-β_4o_ was below the limit of detection. In Dp(16)3Tyb;tgAPP mice in the soluble Triton fraction, amyloid-β_4o_ abundance (F(1,13) = 0.044, p = 0.837) and amyloid-β_42_ abundance (F(1,13) = 0.352, p = 0.563) did not significantly differ from tgAPP littermates at 12-months of age. Dp(16)3Tyb;tgAPP mice (n = 11, 7 male and 4 female) tgAPP (n = 8, 5 male and 3 female). **(C)** In Dp(10)1Yey;tgAPP mice in the soluble Tris fraction, amyloid-β_4o_ abundance (F(1,14) = 0.0003, p = 0.986) and amyloid-β_42_ abundance (F(1,14) = 0.256, p = 0.621) did not significantly differ from tgAPP littermates at 6-months of age. In Dp(10)1Yey;tgAPP mice in the soluble Triton fraction, amyloid-β_4o_ abundance (F(1,14) = 0.147, p = 0.707) and amyloid-β_42_ abundance (F(1,14) = 0.559, p = 0.467) did not significantly differ from tgAPP littermates at 6-months of age. Dp(10)1Yey;tgAPP (n = 8, 3 male and 5 female) tgAPP (n = 12, 7 male and 5 female). In Dp(10)1Yey;tgAPP mice in the soluble Tris fraction, amyloid-β_4o_ abundance (F(1,14) = 1.784, p = 0.203) did not significantly differ from tgAPP littermates at 12-months of age however median amyloid-β_42_ was significantly increased (U(N_Dp(10)1Yey;tgAPP_ = 12, N_tgAPP_ = 8,) = 21, p = 0.039). In Dp(10)1Yey;tgAPP mice in the soluble Triton fraction, amyloid-β_4o_ abundance (F(1,14) = 2.715, p = 0.122) and median amyloid-β_42_ (U(N_Dp(10)1Yey;tgAPP_ = 12, N_tgAPP_ = 8,) = 33, p = 270) did not significantly differ from tgAPP littermates at 12-months of age. Dp(10)1Yey;tgAPP (n = 12, 6 male and 6 female) tgAPP (n = 8, 5 male and 3 female). **(D)** In Dp(17)1Yey;tgAPP mice in the soluble Tris fraction, amyloid-β_42_ abundance (F(1,11) = 0.237 p = 0.636) did not significantly differ from tgAPP littermates at 6-months of age. Amyloid-β_4o_ was below the limit of detection. Dp(17)1Yey;tgAPP (n = 8, 4 male and 4 female) tgAPP (n = 9, 4 male and 5 female). In Dp(17)1Yey;tgAPP mice in the soluble Triton fraction, amyloid-β_4o_ abundance (F(1,11) = 0.490, p = 0.499) and amyloid-β_42_ abundance (F(1,11) = 0.067, p = 0.800) did not significantly differ from tgAPP littermates at 6-months of age. In Dp(17)1Yey;tgAPP mice in the soluble Tris fraction, amyloid-β_42_ abundance (F(1,9) = 0.215 p = 0.654) did not significantly differ from tgAPP littermates at 12-months of age. Amyloid-β_4o_ was below the limit of detection. In Dp(17)1Yey;tgAPP mice in the soluble Triton fraction, amyloid-β_4o_ abundance (F(1,9) 0.58, p = 0.466) and amyloid-β_42_ abundance (F(1,9) = 0.294, p = 0.601) did not significantly differ from tgAPP littermates at 12-months of age. Dp(17)1Yey;tgAPP (n = 8, 3 male and 5 female) tgAPP (n = 7, 4 male and 3 female). Error bars show SEM, data points are independent mice.

**Fig. S6.**
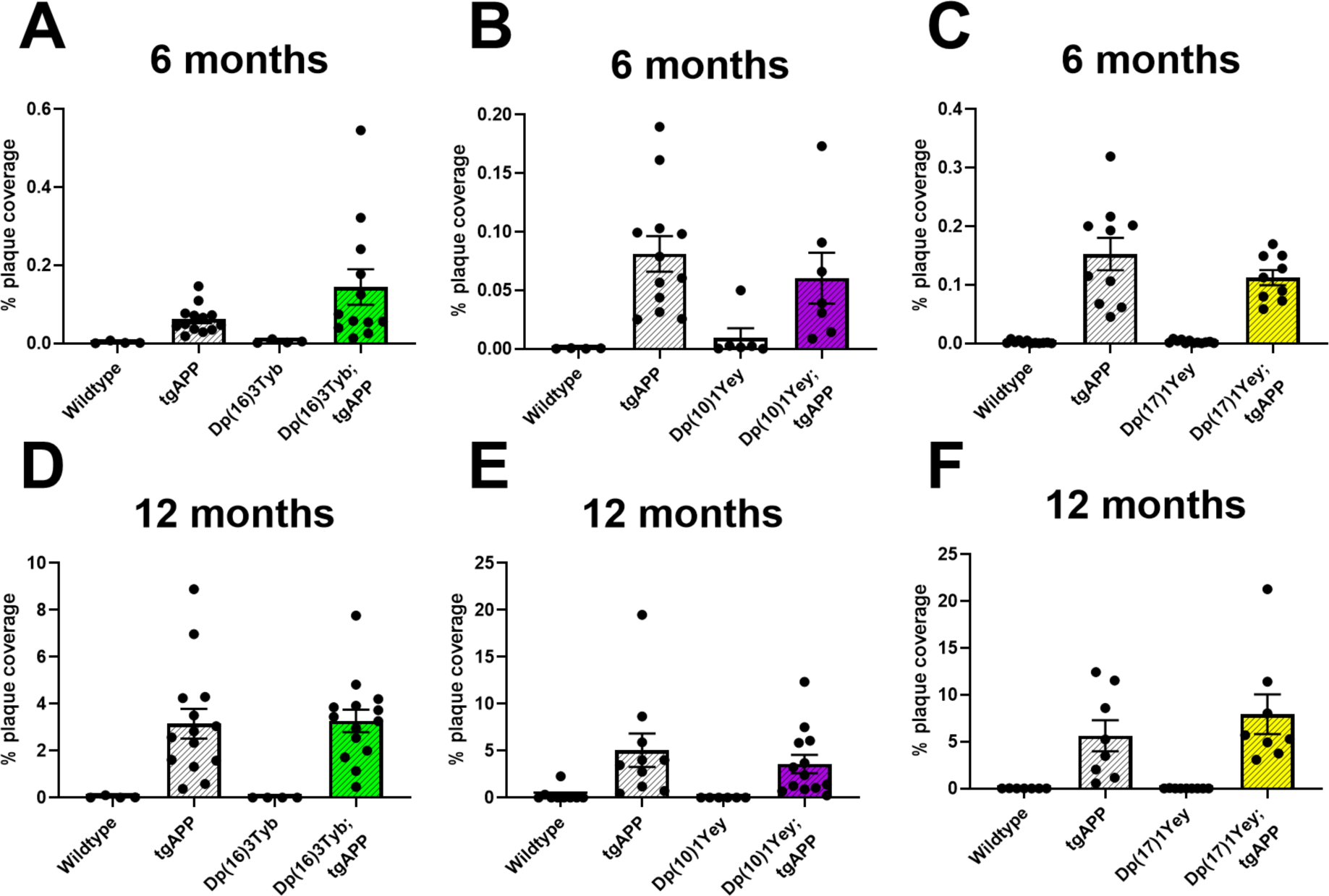
Deposition of amyloid-β in the cortex at 6- and 12-months of age in the Dp(16)3Tyb, Dp(10)1Yey or Dp(17)1Yey tgAPP double mutants. Amyloid-β deposition in the cortex was quantified at (**A-C**) 6- and (**D-F**) 12-months of age in male and female mice, percentage of the region covered by stain was calculated. Error bars show SEM, data points are independent mice. **(A)** No significant difference in amyloid-β deposition in the cortex was detected at 6-months of age in Dp(16)3Tyb;tgAPP compared with tgAPP controls (*U*(N_Dp(16)3Tyb;tgAPP_ = 12, N_tgAPP_ = 13,) = 56, p = 0.247). After systematic outlier testing one tgAPP sample was excluded prior to analysis. Dp(16)3Tyb;tgAPP female n=6, male n=6; tgAPP female n=7, male n=6. **(B)** No significant difference in amyloid-β deposition in the cortex was detected at 6-months of age in Dp(10)1Yey;tgAPP compared with tgAPP controls (F(1,14) = 0.246, p = 0.628). Dp(10)1Yey;tgAPP female n=4, male n=3; tgAPP female n=5, male n=7. **(C)** No significant difference in amyloid-β deposition in the cortex was detected at 6-months of age in Dp(17)1Yey;tgAPP compared with tgAPP controls (*U*(N_Dp(17)1Yey;tgAPP_ = 9, N_tgAPP_ = 10,) = 34, p = 0.400). Dp(17)1Yey;tgAPP female n=5, male n=4; tgAPP female n=6, male n=4. **(D)** No significant difference in amyloid-β deposition in the cortex was detected at 12-months of age in Dp(16)3Tyb;tgAPP compared with tgAPP controls (F(1,23) = 0.031, p = 0.861). Dp(16)3Tyb;tgAPP female n=8, male n=6; tgAPP female n=7, male n=7. **(E)** No significant difference in amyloid-β deposition in the cortex was detected at 12-months of age in Dp(10)1Yey;tgAPP compared with tgAPP controls (F(1,18) = 0.056, p = 0.815). Dp(10)1Yey;tgAPP female n=7, male n=6; tgAPP female n=6, male n=4. **(F)** No significant difference in amyloid-β deposition in the cortex was detected at 12-months of age in Dp(17)1Yey;tgAPP compared with tgAPP controls (F(1,11) = 0.218, p = 0.649). Error bars show SEM, data points are independent mice. Dp(17)1Yey;tgAPP female n=5, male n=3; tgAPP female n=5, male n=3.

